# SYNPLA: A synapse-specific method for identifying learning-induced synaptic plasticity loci

**DOI:** 10.1101/473314

**Authors:** Kim Dore, Yvonne Pao, Jose Soria Lopez, Sage Aronson, Huiqing Zhan, Sanchari Ghosh, Sabina Merrill, Anthony M. Zador, Roberto Malinow, Justus M. Kebschull

**Author notes:** Corresponding authors; Correspondence and requests should be addressed to Roberto Malinow, or Justus Kebschull,.

## Abstract

Which neural circuits undergo synaptic changes when an animal learns? Although it is widely accepted that changes in synaptic strength underlie many forms of learning and memory, it remains challenging to connect changes in synaptic strength at specific neural pathways to specific behaviors and memories. Here we introduce SYNPLA (SYNaptic Proximity Ligation Assay), a synapse-specific, high-throughput and potentially brain-wide method capable of detecting circuit-specific learning-induced synaptic plasticity.

---

Changes in synaptic strength have been hypothesized to underlie learning and memory since before the time of Hebb. The best studied form of Hebbian synaptic plasticity is long-term potentiation (LTP), which underlies the formation of fear conditioning memories^1^. Work over the last two decades has uncovered the molecular mechanisms of LTP in great detail, revealing that the expression of plasticity is mediated by the rapid synaptic insertion of GluA1-subunit containing AMPA receptors mobilized from an extrasynaptic pool^2^. These GluA1-containing receptors are then likely replaced by GluA1-lacking AMPA receptors within 15 to 72 hours^3–5^, making synaptic GluA1 a marker of recently potentiated synapses. LTP appears to be very general at glutamatergic synapses in the central nervous system, including in cortex^3^, amygdala^6^, ventral tegmental area^7^, nucleus accumbens^8^, striatum^9^ and lateral habenula^10^. Furthermore, plasticity mediated by trafficking of GluA1-subunit containing AMPA receptors appears to play an important part in several neuropsychiatric disorders, such as addiction^11^, autism^12^ and depression^10^.

Relating behaviorally-induced plasticity to its synaptic substrate remains an important technical challenge. Electrophysiological or optical methods provide mechanistic insight, but are low-throughput and require specialized equipment^6^. Somatic expression of immediate early genes (IEG) such as *cfos* can be used to screen for brain areas activated during plasticity^13^, but lack synaptic resolution. We therefore sought to develop a pathway-specific histological method for detecting behaviorally induced changes.

Here we present SYNPLA, a method that uses the proximity ligation assay (PLA) to detect synaptic insertion of GluA1-containing AMPA receptors in defined circuits. PLA is a highly sensitive and specific biochemical method that reliably detects the close (< 40 nm) juxtaposition of two proteins *in situ*^14^ (Fig. 1**a**). To detect nearby molecules A and B, PLA uses a standard primary antibody raised against A, and another raised against B. These antibodies are detected by secondary antibodies conjugated to unique oligonucleotides (Ao, Bo). A second set of oligonucleotides, ABo1 and ABo2, that are complementary to parts of both Ao and Bo are added. Only when Ao and Bo are sufficiently close can ABo1 and ABo2 be ligated to form a circle. This circle is then amplified (>1,000-fold) via rolling-circle amplification to form a nanoball of DNA, which can be reliably probed with complementary fluorescent nucleotides and observed as a punctum with light microscopy. A key advantage of PLA over traditional immunostaining is the absolute requirement for a pair of proteins to produce a PLA signal—a logical AND—which confers high specificity and thus reduces the false positive rate. Moreover, the 1000-fold signal amplification renders each positive signal very bright and therefore easily distinguishable from background, allowing results to be imaged quickly, and at lower resolution than would otherwise be necessary for resolving synaptic contacts.

**Figure 1:**
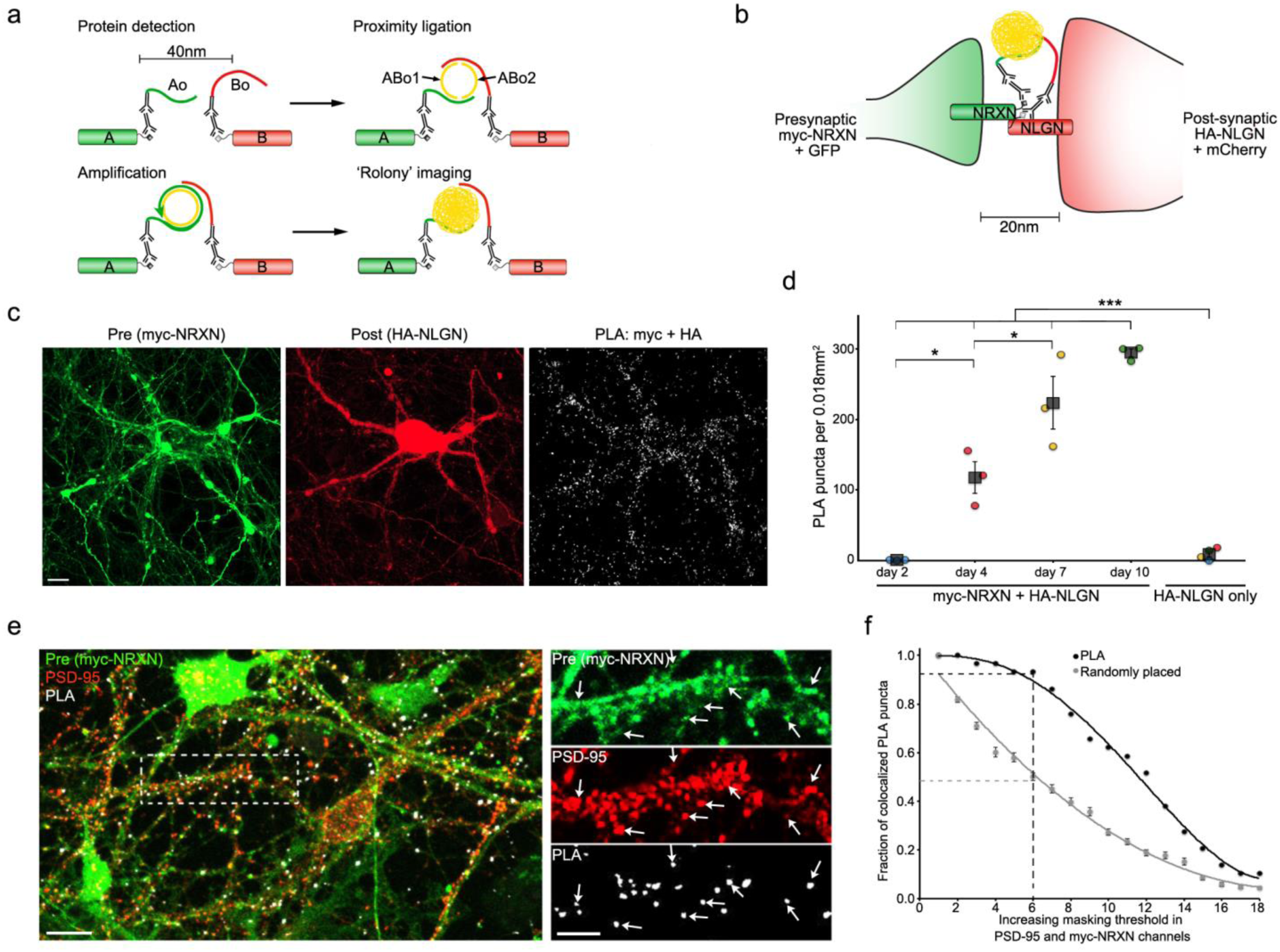
SYNPLA detects synapse formation and labels synapses. **a**) Diagram of a general PLA reaction. Note that A and B must be sufficiently close to permit complementary binding and ligation of ABo1 and ABo2 and subsequent amplification to make a rolony. **b**) Diagram of PLA targeting recombinant presynaptic myc-NRXN and postsynaptic HA-NLGN. **c**) PLA rolonies (white dots, right) formed between postsynaptic cultured mouse hippocampal neuron expressing HA-NLGN + mCherry (red; middle) and co-cultured presynaptic neurons expressing myc-NRXN + GFP (green; left). Scale bar: 10 μm. **d**) PLA puncta in each sample (circles) and average (squares) ± SEM for indicated expression and days *in vitro* (color indicates data acquired on the same day). *** p<0.001, 2-way ANOVA (see Methods); * p<0.05, 1-way ANOVA (with Tukey-Kramer post-hoc test). **e**) PLA reaction between endogenous GluA1 and recombinant myc in cultured rat hippocampal neurons (exposed to cLTP, see Figure 2) expressing myc-NRXN and cytosolic fluorescent protein (green), subsequently immunostained for endogenous PSD-95 (red). Scale bars: 10 μm (left); 5 μm (right). **f**) PLA puncta localize to synapses. In PSD-95 and myc-NRXN channels (from **e**, right), pixels were set to zero (masked) for values below a progressively increasing threshold (x axis). Puncta (PLA or randomly placed) display colocalization if non-zero pixels exist within 0.14 μm in both thresholded channels. For indicated threshold, ∼95% of PLA, while only ∼50% of randomly placed (mean ± SEM of 30 placements) puncta colocalized with pre- and post-synaptic markers. See Supplementary Fig. 3 and Methods for details.

We first confirmed previous experiments^15^ suggesting that PLA could be used to detect direct protein-protein interactions across the synaptic cleft in dissociated cultured neurons (Fig. 1**b-d**). We exploited the developmental window (days 2-10) during which synapses are formed in this preparation^16^. In some neurons we expressed the presynaptic protein Neurexin1b with a myc tag (myc-NRXN) along with cytosolic GFP; in other neurons we expressed Neuroligin1, the normal postsynaptic binding partner of NRXN^17^, with an HA tag (HA-NLGN) and cytosolic mCherry. Consistent with the normal time course of synapse formation^16^, PLA reactions using antibodies to myc and HA produced an increasing number of PLA puncta, whereas cultures expressing only HA-NLGN led to minimal PLA products highlighting the specificity and high signal to noise of SYNPLA (Fig. 1**c-d**, Supplementary Figs. 1 and 2). To demonstrate that SYNPLA labels synapses, we measured colocalization between PLA puncta (generated as described below) and both presynaptic signal and post-synaptic PSD-95 (visualized with immunofluorescence; see Fig. **1e-f** and Supplementary Fig. 3 for details). We also performed PLA using antibodies to endogenous pre- and postsynaptic proteins (Supplementary Fig. 4), showing protein overexpression is not required. These results indicate that SYNPLA can detect synapses expressing recombinant and/or endogenous proteins with a high signal to noise ratio.

We reasoned that since the 20 nm synaptic cleft is less than the 40 nm PLA capture radius, SYNPLA could detect selectively the apposition of presynaptic NRXN with GluA1 inserted into the postsynapse during LTP. In contrast, no PLA products should be produced between NRXN and the more distant extrasynaptic pool of uninserted GluA1-containing AMPA receptors (Fig. 2**a**). We expressed myc-NRXN in dissociated cultured neurons (Fig. 1**e-f;** Fig. 2**b-d**) or in region CA3 of cultured hippocampal brain slices^18^ (Fig. 2**e-g**), and we performed PLA using antibodies to myc and the extracellular domain of GluA1 on the dissociated cultured neurons or region CA1 of organotypic slices. Although we observed few PLA puncta in control conditions, chemically induced LTP (cLTP)^19^ greatly increased the number PLA puncta (Fig. 2**b-g**; 4-fold, p<0.01, paired t-test in dissociated cultures; 12-fold, p<0.001, paired t-test in organotypic cultures). The increase in the number of PLA puncta was blocked by addition of APV, a blocker of LTP induction^20^, to neurons prior to cLTP (Fig. 2**b-d**; p<0.01, unpaired t-test). These results indicate that SYNPLA can employ recombinantly expressed presynaptic myc-NRXN and endogenous postsynaptic GluA1 to detect synaptic plasticity under cultured conditions.

**Figure 2:**
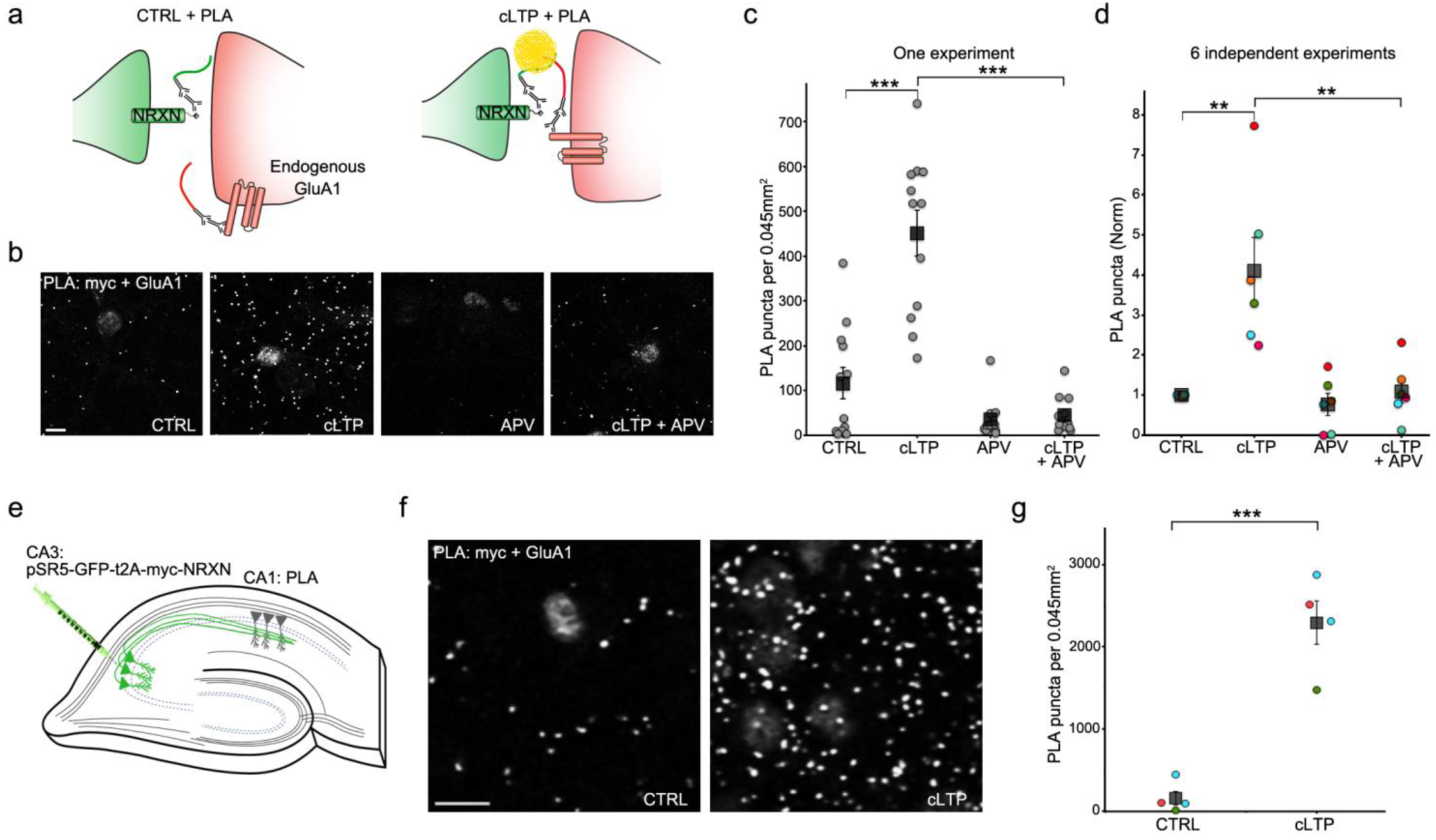
SYNPLA detects synaptic potentiation in hippocampal primary cultures and organotypic slices. **a**) Diagram of SYNPLA targeting presynaptic myc-NRXN and. cLTP induces mobilization of GluA1 from an extrasynaptic pool and insertion into the post-synaptic density, decreasing the distance between the targeted proteins and permitting PLA. **b**) Representative images of SYNPLA reactions performed on myc-NRXN expressing cultured rat hippocampal neurons at 14 days *in vitro* for the indicated conditions; PLA shown in gray. Scale bar: 10 μm. **c**) Quantification of a single SYNPLA experiment (as in **f**). The number of SYNPLA puncta detected in each field of view (round) and the average (square) ± SEM across all fields of view are shown; *** p<0.001, 1-way ANOVA followed by Tukey-Kramer post-hoc test. **d**) Quantification of six independent experiments (indicated by different colors); the average number of SYNPLA puncta across 10-12 fields of view for each experiment (circles, normalized to CTRL) and the average PLA signal across experiments (square) ± SEM are shown; ** p<0.01, 1-way ANOVA (with Tukey-Kramer post-hoc test). **e**) Diagram of SYNPLA in rat organotypic hippocampal slices. Sindbis virus expressing myc-NRXN is injected in the presynaptic CA3 region SYNPLA between myc-NRXN and endogenous GluA1 is measured in the postsynaptic CA1 region. **f**) Representative images of SYNPLA under the indicated conditions in region CA1 of organotypic slice cultures. Scale bar: 10 μm. **g**) Quantification of SYNPLA puncta in organotypic slices, from three independent experiments (indicated with different colors, 4 slices per condition (one circle per slice)); squares indicate average ± SEM across experiments; *** p<0.001, paired t-test.

To test if SYNPLA can detect synaptic plasticity following memory formation *in vivo*, we injected the auditory cortex and/or the medial geniculate nucleus of the thalamus of rats with a virus expressing myc-NRXN and cytosolic GFP. We then exposed the rats to a cued fear-paired conditioning protocol, wherein a 10 second tone (conditioned stimulus) is immediately followed by a brief foot shock (Fig. 3**a-c**). Such protocols have been shown to produce LTP-like plasticity at synapses onto the lateral amygdala (LA)^6^. Control animals received either no viral injection, no conditioning or unpaired conditioning, wherein the tone and the foot shock are not temporally paired. We perfused control or conditioned animals 30 minutes after conditioning, and then post-fixed and sectioned the brains at 50 μm. We subjected tissue sections to SYNPLA, and imaged the LA region containing GFP labeled presynaptic fibers (Fig. 3**d**). We found that uninjected, naïve or animals receiving unpaired conditioning displayed few PLA puncta, whereas animals receiving paired conditioning displayed on average three-fold increase in PLA puncta (Fig. 3**f-h**; p<0.001, paired t-test; see also Supplementary Figs. 6 and 7). The large fractional increase in GluA1-containing receptors at synapses during learning is consistent with a synaptic plasticity model wherein GluA1-containing receptors are added during plasticity and replaced within about 24 hours with GluA1-lacking receptors^3^. These results indicate that SYNPLA can detect synaptic plasticity induced by learning.

**Figure 3:**
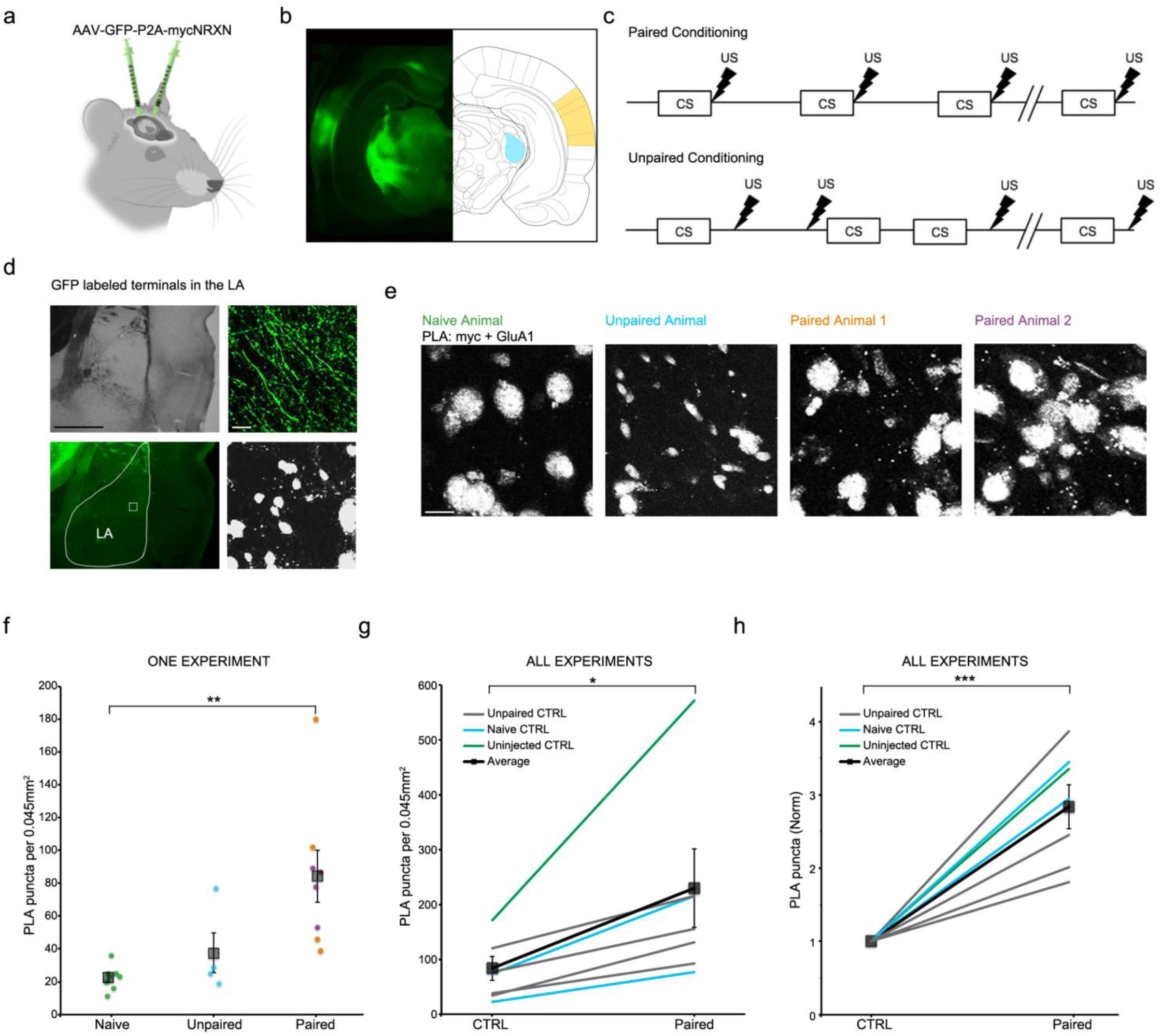
SYNPLA detects potentiated synapses between medial geniculate nucleus or auditory cortex and the lateral amygdala (LA) following fear conditioning. **a**) Injection of AAV9-GFP:P2A:myc-NRXN into auditory cortex and/or medial geniculate nucleus. **b**) GFP expression in the cell bodies at the injection sites (Auditory cortex: yellow, medial geniculate nucleus, blue). **c**) Diagram of paired (top) and unpaired (bottom) fear conditioning paradigm; tone (CS; 10 seconds) and shock (thunderbolt) delivered where indicated. **d**) Representative images of presynaptic GFP-labeled fibers terminating onto the lateral amygdala (LA) under low (left; top: transmitted light; below: GFP signal, LA traced in white) and high (right; top: GFP signal; PLA shown below) magnification in an animal that received paired conditioning. Scale bar: left, 1 mm; right, 10 μm. **e**) Representative images of a single SYNPLA experiment in the LA from animals subjected to the indicated conditions; colors correspond to symbols in **f**. Scale bar: 10 μm. **f**) Quantification of the experiment shown in **e**. Number of SYNPLA puncta detected in one field of view (circles) and average across all fields of view (squares) ± SEM; * p<0.05, ** p<0.01; unpaired t-test. **g**) Quantification of all *in vivo* experiments, showing SYNPLA puncta per animal (average across all fields of view; gray) under the indicated control conditions and the average across animals (square) ± SEM are shown; * p<0.05; paired t-test. See Supplementary Fig. 6 for information regarding specific injection sites used in these experiments. Supplementary Fig. 7 shows similar results when data are normalized to GFP labeled fibers intensity. **h**) Same data as in **g**, normalized to control, ** p<0.01; paired t-test.

Here we have demonstrated that SYNPLA can reliably detect synaptic potentiation in circuits participating in newly formed memories. SYNPLA can be used to probe plasticity in anatomically defined pathways as in the present study, or in genetically defined pathways by limiting expression of myc-NRXN only to neurons expressing Cre recombinase. Simultaneous screening of multiple pathways in the same animal might also be possible by exploiting orthogonal PLA probes^21^. Unlike IEG-based screening approaches, which assay cell-wide changes, SYNPLA is synapse- and pathway-specific, and directly assesses synaptic plasticity. Moreover, it is fast and easy to perform, and so could be scaled up as a powerful brain-wide screen for behaviorally-induced plasticity.

## Materials and Methods

### Experimental model

#### Primary neuronal cultures

For the time course of interacting proteins shown in Fig. 1**c-d**, we obtained single cell suspension of hippocampal neurons from E18 CD1 (Charles River) mouse embryos as previously described^22^. We separately nucleofected these neurons with CAGs:gfp-P2A-myc-Nrxn1B (jk95; available on Addgene with accession 120176) or CAGs:mcherry-P2A-HA-Nlgn1 (jk96; available on Addgene with accession 120177) using a Lonza Nucleofector and Mouse Neuron transfection kit (Lonza) according to the manufacturer’s instructions. We then either mixed cells nucleofected with CAGs:gfp-P2A-myc-Nrxn1B or CAGs:mcherry-P2A-HA-Nlgn1 and co-plated them, or plated cells nucleofected with CAGs:mcherry-P2A-HA-Nlgn1 only as a negative control. We fixed samples from the same cell suspension (thus controlling for variations in cell density) in 4% PFA at 2, 4, 7 and 10 days *in vitro* (DIV) for 15 min at room temperature and performed PLA without delay.

For experiments shown in Fig. 1**e-f** and 2**b-d**, we prepared primary rat hippocampal neurons from postnatal day 0-1 rat pups as previously described^23^. At 12-15 DIV, we infected the neurons with a Sindbis virus expressing mCherry-t2A-myc-Nrxn1B. Eighteen to 24 hours later, we subjected neurons to cLTP for 10 min (in HBSS based solution, 100 nM rolipram, 50 μM forskolin, 100 μM picrotoxin, 2 mM Ca^2+^, 0 mM Mg^2+^)^24^. Control neurons were kept in HBSS for 10 min. When indicated, 100 μM APV was added. Immediately after cLTP, we fixed the neurons in 4 % PFA for 10 min at room temperature and performed PLA without delay.

#### Organotypic hippocampal slices

For experiments shown in Fig. 2**e-g**, we prepared organotypic hippocampal slices from postnatal day 7-8 rat pups as previously described^18^. Slice cultures were infected in the CA3 region with the mCherry-t2A-myc-Nrxn1B Sindbis at 14–18 DIV. Eighteen to 24 hours later, we subjected the slices to cLTP or mock stimulation similar as with the primary rat hippocampal cultures. Briefly, we incubated the slices in ACSF+drugs (118 mM NaCl, 2.5 mM KCl, 26 mM NaHCO_3_, 1 mM NaH2PO4, 20 mM glucose, 0 mM MgCl_2_, 2 mM CaCl_2_, 100 nM rolipram, 50 µM forskolin, 100 µM picrotoxin) or regular ACSF (118 mM NaCl, 2.5 mM KCl, 26 mM NaHCO_3_, 1 mM NaH_2_PO_4_, 1 mM MgCl_2_, 2 mM CaCl_2_, 20 mM glucose) for 16 min, and immediately fixed the slices in 4 % PFA for 4 hours at 4C. To make sure that this cLTP protocol produced functional synaptic potentiation, we performed evoked field recordings in 4 independent slices (see Supplementary Figure 5). For electrophysiology, fEPSPs were recorded until 20 minutes of a stable baseline was acquired. Following the stable baseline, ACSF was washed out and replaced with cLTP ACSF for 16 minutes, during which time no electrical stimulation was applied. The cLTP solution was subsequently washed out and replaced with ACSF as described in the previous section, and electrical stimulation was resumed at a frequency of 0.1 Hz.

#### Animals for ex vivo SYNPLA

We injected 6- to 8-week-old WT Sprague-Dawley rats in the medial geniculate nucleus (AP: −6.4mm, ML: – 3.4 mm, DV: 5.4 mm) and/or the auditory cortex (AP: −5mm, ML: 7.2 mm, DV: 4.5 mm) with 500 nl of AAV9-GFP-P2A-myc-Nrxn1B (construct jk95; UNC Vector Core). See Supplementary Fig. 6 for details on the different injection sites for all animals tested. Virus was injected over a 4-minute period and the injection pipette was kept in place for an additional 10 minutes to ensure focal infection. After a minimum of 4 weeks, injected animals were subjected to paired or unpaired tone-conditioning paradigm (Fig. 3**c**). Briefly, in the paired condition, a 10 second tone (1.2 kHz, 70 dB) was followed immediately by a 1 second footshock (0.8 mA delivered via Coulbourn Precision Animal Shocker) with an inter-trial interval of 60 seconds with a 20 second jitter. In the unpaired condition, the animal was on the same shock schedule but the tones were played at pseudorandom intervals such that they did not predict the shock. This behavioral paradigm was programmed via a custom MATLAB script and connected to the Shocker via a National Instruments DAQ. Thirty minutes after conditioning, we anaesthetized the animals with isoflurane and transcardially perfused them with 4 % PFA. We postfixed the brains for 36 hours at 4C, and then transferred them to PBS before cutting 50 µm coronal sections on a vibratome. We performed PLA on the tissue section where the lateral amygdala (LA) was most visible (as shown in Fig. 3**d**). In total, we performed 6 independent *ex vivo* experiments. Five of these experiments consisted of one control animal (uninjected, naïve or unpaired conditioning) and one animal that received paired conditioning. In one experiment (shown in Fig. 3**e-f**), however, we processed 4 animals (2 controls and 2 paired) at the same time. For summary graphs shown in Fig. 3**g-h**, each control animal of this experiment was randomly paired with one animal from the same experiment that received paired conditioning.

Animal procedures were approved by the Cold Spring Harbor Laboratory or UCSD Animal Care and Use Committee and were carried out in accordance with National Institutes of Health standards.

### Proximity ligation assay (PLA)

We performed PLA using Duolink PLA reagents (Sigma Aldrich) largely according to the manufacturer’s instructions. For experiments on primary neuronal cultures, we first incubated the samples in 4 mM glycine for 5 min. Organotypic or *ex vivo* slices were permeabilized for 30 min using 0.2 % TritonX-100. We then blocked all samples using Duolink blocking buffer (organotypic slices were blocked with BlockOneHisto (Nacalai Tesque)) at 37C for 1 hour. We then applied primary antibodies (for Fig. 1**c-d**: goat anti-myc (ab9132, Abcam) and rabbit anti-HA (ab9110, Abcam), both at 1:10,000 dilution; Fig. 1**e-f** and Figs. 2 and 3: goat anti-myc (ab9132, Abcam), 1:2000 dilution and rabbit anti-GluA1 (AGC-004, Alomone labs), 1:100 dilution (1:50 for Fig. 3)) in Duolink antibody diluent for 1 hour at room temperature or overnight at 4C. For Supplementary Fig. 2 different primary antibodies were used as negative controls: guinea pig anti-GluA1 (AGP-009, Alomone labs), rabbit anti-cFos (Cell signaling, #2250); all 1:100 dilution. For Supplementary Fig. 4 an antibody targeting endogenous NRXN (goat polyclonal from abcam, ab77596) was used (1:100 dilution) instead of the anti-myc antibody. After washing 3Ç10 min in PLA wash buffer A, we probed the primary antibodies using Duolink PLA probes Goat PLUS and rabbit MINUS at 37C for 2 hours, and then used Duolink in Situ Detection Reagents FarRed (Sigma Aldrich) to detect proximity between the proteins of interest.

### Imaging

For Fig. 1**c-d**, we subjected 3 separate coverslips containing both myc-NRXN and HA-NLGN expressing neurons, as well as one coverslip containing only HA-NLGN expressing neurons to PLA at each time point. Per coverslip, we acquired 3 z-stacks containing a single mCherry expressing cell using a 63x oil objective on a Zeiss laser scanning 780 confocal.

For Fig. 2**b-c**, we acquired 12 z-stacks per condition (1 coverslip per condition). In Fig. 2**d**, each dot is the average number of PLA puncta in 10-12 images of a single experiment, normalized to its control sample for each of 6 independent experiments. For Fig. 2**f-g**, we used 1-2 slices per condition for each experiment. Three independent repeats are shown, totaling 4 slices per condition.

For Fig. 3, we acquired 2-5 z-stacks in the LA of each slice (one slice per animal). Fig. 3**f** shows the number of PLA puncta detected for each z-stack for the indicated animals. In Fig. 3**g**, each dot represents the average number of PLA puncta for all individual animals. Fig. 3**h** shows the same data as 3**g** but the average PLA signal in each paired animal was normalized to its control.

All images used for experiments shown in Fig. 1**e-f** and Figs. 2 and 3 were acquired on an Olympus FV1000 confocal microscope with a 60x oil-immersion objective. All images were obtained blind to condition.

### Analysis and statistics

#### PLA puncta detection

For all experiments in primary cultures, we quantified the number of PLA dots across the entire thickness of the cell layer using a custom Fiji macro (consisting primarily of a maxima finding function). For organotypic slices samples, we quantified the number of PLA puncta in the CA1 region using a custom Fiji macro in a single z-plane, approximately 2-3 μm below the slice surface. PLA signal decreases to noise levels deeper into the tissue (Supplementary Fig. 5), likely due to limited reagent penetration. For *ex vivo* experiments, we quantified PLA across the entire thickness of the samples to account for tissue irregularity with a modified Fiji macro that included a nuclei subtraction step. All images were analyzed blind to condition.

#### Colocalization analysis

For experiments shown in Fig. 1**e-f**, we expressed myc-NRXN/cytosolic mCherry in 14 DIV hippocampal neurons then induced cLTP (as described in the Primary neuronal cultures section) in order to deliver endogenous GluA1 to synapses. Neurons were fixed and PLA between myc-NRXN and endogenous GluA1 was performed. At the same time, we also immunolabeled for PSD-95 (mouse monoclonal antibody MA1-045, ThermoFisher), a well-established synaptic marker. To determine if PLA signal colocalizes with PSD-95, we conducted the following analytical process, performed with a MATLAB script. For each image, we identified the location of PLA puncta in a given region of interest. Then we progressively increased the masking threshold for the presynaptic fiber (mCherry) and postsynaptic (PSD-95 immunolabel) channels, setting image values below this threshold to zero. For each identified PLA punctum, and for each masking level, we quantified the number of non-zero pixels within 0.14 μm (1 pixel) of the PLA punctum in the presynaptic AND postsynaptic image. We then repeated this analysis 30 times, but using randomly placed puncta instead of PLA signals. See Fig. 1**e-f** and Supplementary Fig. 3 for more details.

#### Statistics

For experiments shown in Fig. 1**d**, we tested for statistical significance of increased PLA puncta when both epitopes were present using a 2-way ANOVA (using DIV, expressed epitopes (myc and HA or HA only) as factors). We used a 1-way ANOVA and Tukey-Kramer post-hoc testing to assess significance of PLA puncta increase over DIV. For other experiments where we compared multiple conditions (Fig. 2**c-d** and Fig. 3**f**) we used a 1-way ANOVA and Tukey-Kramer post-hoc testing to assess significance. Unpaired or paired t-test were used elsewhere as indicated.

## End Matter

### Author Contributions and Notes

Experiments were performed and analyzed by K.D., Y.P., J.S.P., and S.A., (Figs 1e-f, 2, 3; Supplementary Figs. 2-7; R.M. laboratory), and H.Z., S.G., S.M., and J.M.K. (Fig 1a-d; Supplementary Fig 1; A.M.Z. laboratory). K.D., A.M.Z, R.M. and J.M.K wrote the manuscript and supervised the project.

This article contains supporting information online.

### Competing interests

The authors declare no competing financial interests.

## Acknowledgments

Funding sources: National Institutes of Health (5RO1NS073129 and 5RO1DA036913, A.M.Z.; 29R01MH049159, R.M.; U01MH109113, A.M.Z and R.M.); Brain Research Foundation (BRF-SIA-2014-03, A.M.Z.); IARPA (MICrONS D16PC0008, A.M.Z.); Simons Foundation (382793/SIMONS, A.M.Z.); Paul Allen Distinguished Investigator Award (A.M.Z.); Shiley Foundation (R.M.); Boehringer Ingelheim Fonds PhD fellowship (J.M.K.); Genentech Foundation PhD fellowship (J.M.K).

We would also like to thank the Finkelstein Lab for providing the word Biorxiv-Template used to typeset this article.

