## Supplemental Figures for "SYNPLA: A synapse-specific method for identifying learning-induced synaptic plasticity loci"

* Corresponding authors

### Supplementary Figures

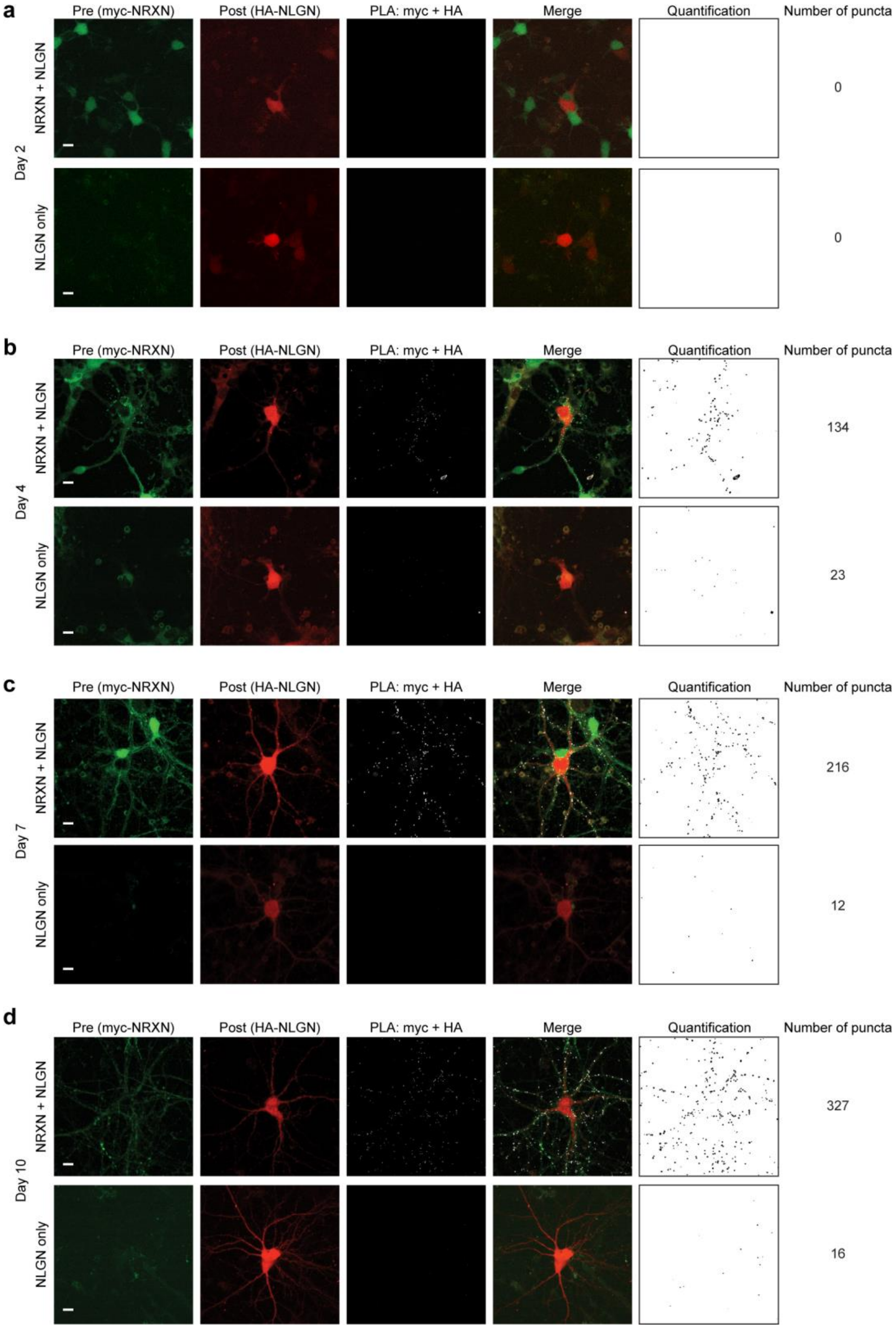


**Supplementary Figure 1 (related to Figure 1b-d): SYNPLA between myc-NRXN and HA-NLGN across time in culture**

Representative images of PLA signals (3rd column) generated by probing for HA and myc epitopes in co-cultures of HA-NLGN+mCherry expressing neurons neurons (2nd column) and myc-NRXN+GFP expressing neurons; 1st column), or in control cultures of only HA-NLGN+mCherry expressing neurons, after different days in culture. **a**) 2 days in vitro, **b**) 4 days in vitro, **c**) 7 days in vitro, **d**) 10 days in vitro. Automatically detected PLA puncta are shown in the last column, together with the number of puncta detected in each image. Scale bar is 10 μm.

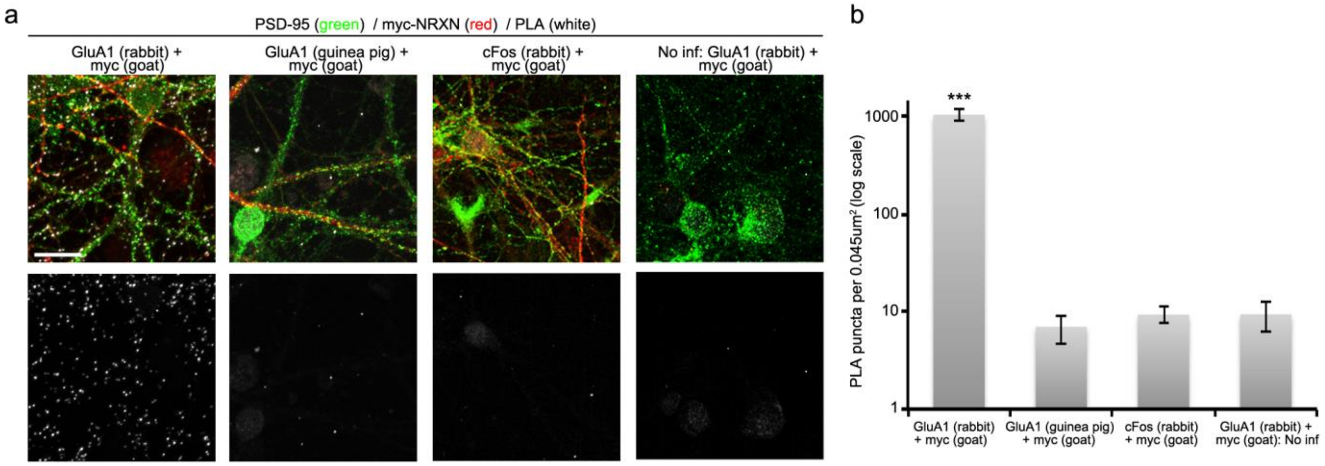


**Supplementary Figure 2 (related to Figures 1 and 2) SYNPLA displays high specificity and very low background**

**a)** Example images of SYNPLA in 15 DIV cultured neurons, probed with different primary antibodies; target, and animal used to raise antibody indicated; PLA probes used are always anti-rabbit and anti-goat. Top: Composite images showing PSD-95 staining in green, cytosol of myc-NRXN-expressing neuron in red and PLA in white. Images shown below displays PLA signal only. Scale bar is 20μm.

**b)** Quantification of SYNPLA puncta in the indicated conditions, a log scale is used for better data visualization. Note that all negative controls yield ~100-fold less PLA signal.

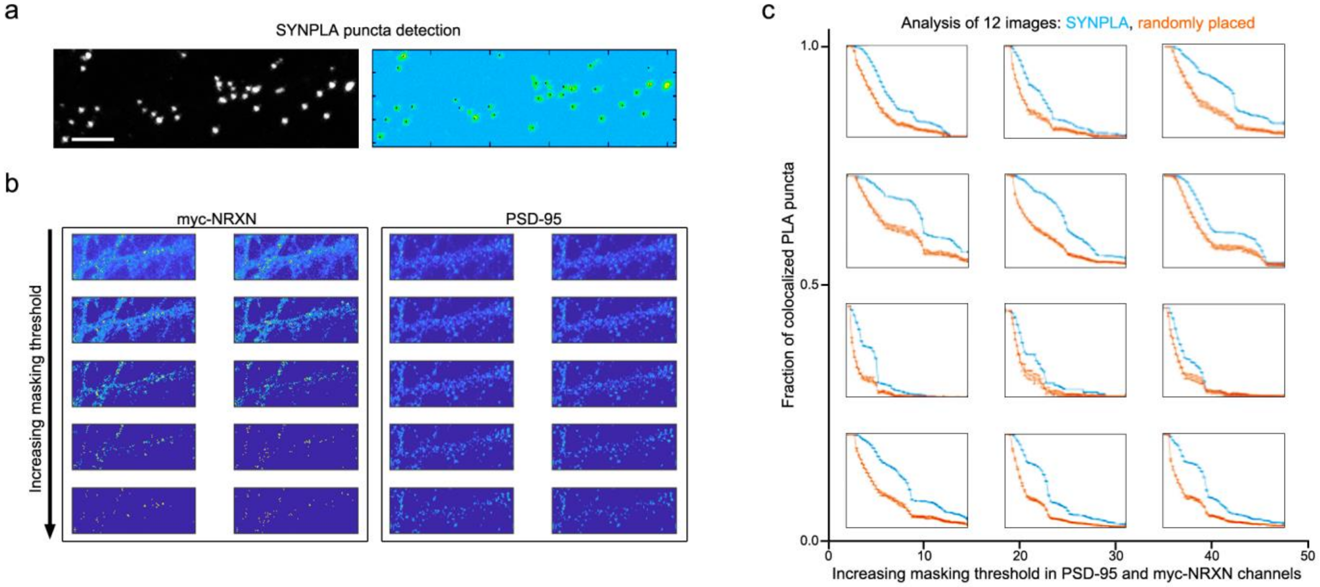


**Supplementary Figure 3 (related to Figure 1e-f): Approach to detect colocalization of SYNPLA signal with pre- and post-synaptic signals**

**a)** Automatic detection of SYNPLA puncta. Scale bar is 5μm.

**b)** Increasing masking threshold on presynaptic signal (myc-NRXN) and postsynaptic signal (PSD-95).

**c)** Result of colocalization analysis on 12 different images, SYNPLA shown in blue and randomly placed puncta (30 trials) shown in orange.

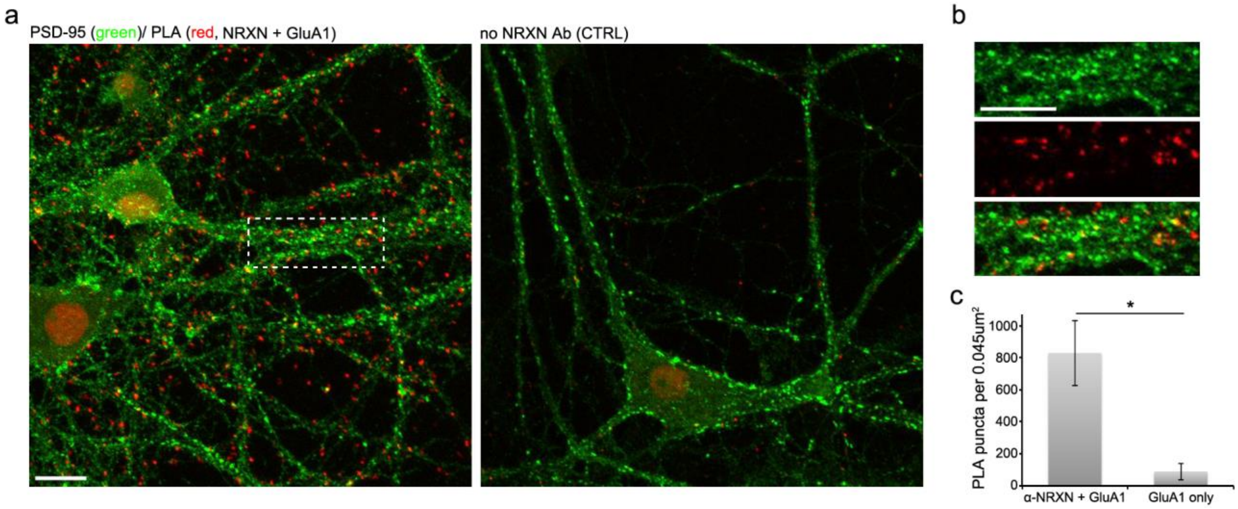


**Supplementary Figure 4: SYNPLA between endogenous Neurexin and GluA1**

SYNPLA in cultured hippocampal neurons (after cLTP) between endogenous NRXN and GluA1.

**a)** Example images of endogenous SYNPLA (left) or negative control where the NRXN antibody was omitted (right). Synaptic labeling is demonstrated by a PSD-95 co-staining in green. Scale bar is 10μm.

**b)** Inset of SYNPLA shown in **a)** with separated channels. Scale bar is 10μm.

**c)** Quantification of SYNPLA puncta in the indicated conditions.

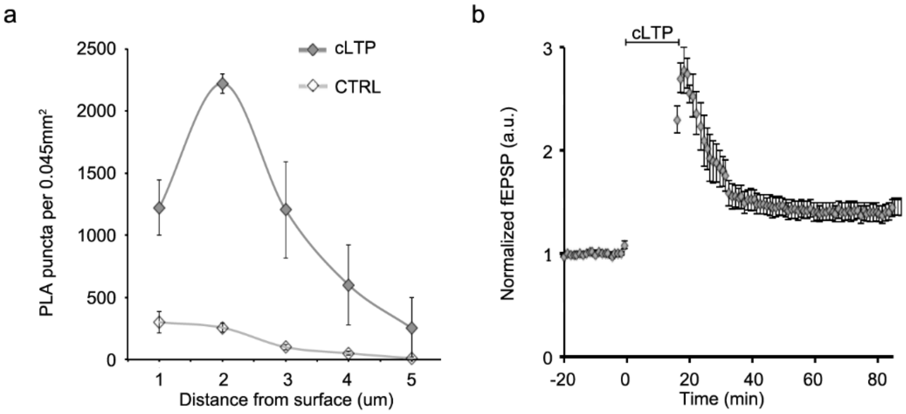


**Supplementary Figure 5 (related to Figure 2e-g): SYNPLA between myc-NRXN and endogenous GluA1 in organotypic slices**

**a**) Number of PLA puncta quantified in each 1um z-slice in one representative experiment in hippocampal organotypic slices, 2 slices (2 z-stacks per slice) in each condition.
**b**) Evoked field potential recordings were performed in organotypic slices subjected to the cLTP protocol, which led to robust potentiation, n=4 slices.

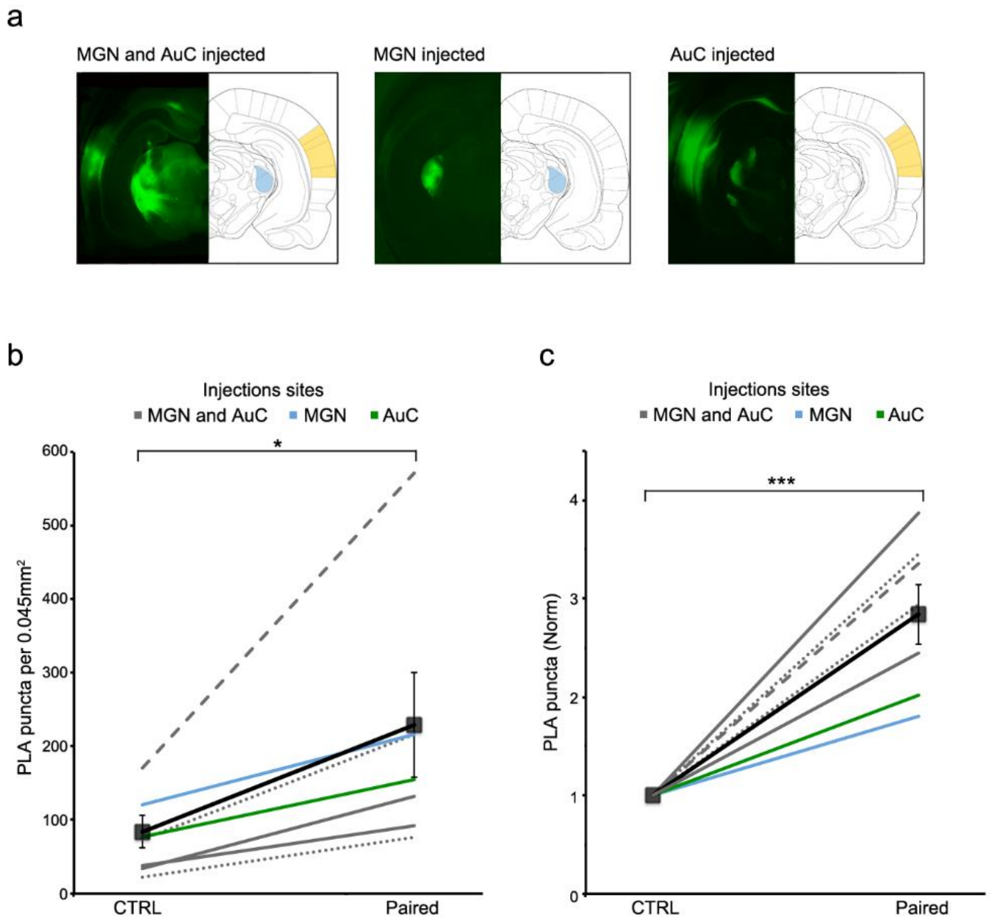


**Supplementary Figure 6 (related to Figure 3): Injection sites used for *ex-vivo* SYNPLA**

**a**) Representative images of the injection sites used in this study.
**b**) Quantification of all *ex vivo* experiments; SYNPLA puncta per animal (average across all fields of view) under the indicated conditions (CTRL: uninjected (dashed line) naïve (dotted lines) or unpaired (solid lines)) and the average across animals (square) ± SEM are shown; * $p<0.05$; paired t-test. Animals that were injected in the medial geniculate nucleus (MGN) and auditory cortex (AuC) are shown in gray; animals injected in the MGN only are shown in blue and animals injected in the AuC only are shown in green.
**c**) Same data as in **b**, normalized to control, *** $p<0.01$; paired t-test.

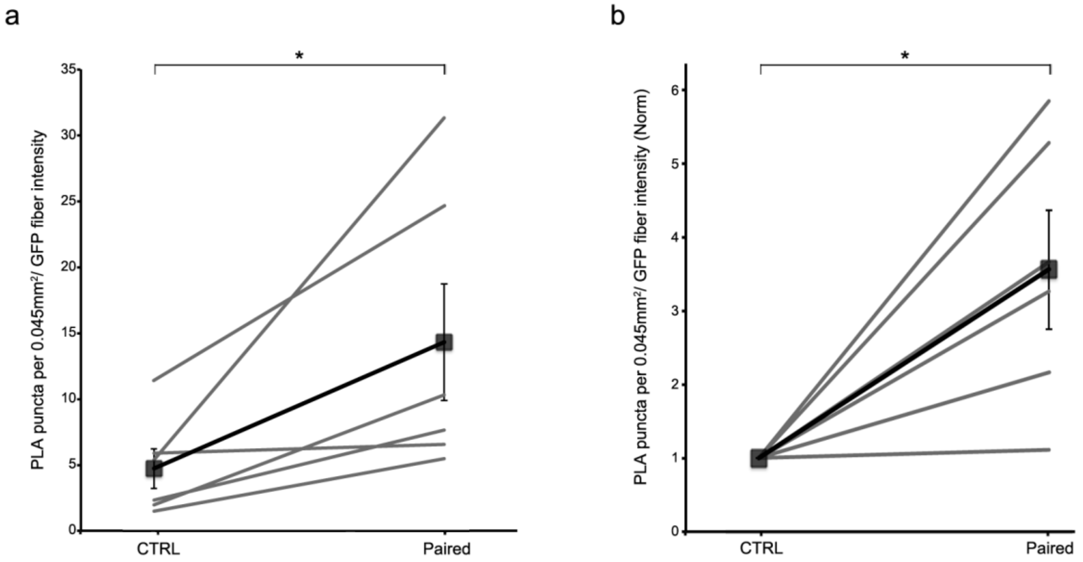


**Supplementary Figure 7 (related to Figure 3):** ***Ex vivo*** **data normalized to myc-NRXN green fiber intensity**

**a**) Quantification of 6 of the 7 *ex vivo* experiments (one experiment was removed because the control animal was uninjected); number of SYNPLA puncta per animal (average across all fields of view) normalized to the pre-synaptic GFP fiber intensity in the same field of view. Each experiment is shown as a gray line and the average across animals is shown in black (square) ± SEM are shown; * $p<0.05$; paired t-test.

**b**) Same data as in **a**, normalized to control, * $p<0.05$; paired t-test.
